# Platelet miRNA bio-signature discriminates between dementia with Lewy bodies and Alzheimer disease

**DOI:** 10.1101/2020.05.04.075713

**Authors:** Ana Gámez-Valero, Jaume Campdelacreu, Dolores Vilas, Lourdes Ispierto, Daniela Samaniego, Jordi Gascón-Bayarri, Ramón Reñé, Ramiro Álvarez, Maria P Armengol, Francesc E. Borràs, Katrin Beyer

**Affiliations:** Department of Pathology, Germans Trias i Pujol Research Institute (IGTP), Universitat Autònoma de Barcelona (UAB), Barcelona, Spain; REMAR-IVECAT group, Germans Trias i Pujol Research Institute (IGTP), Badalona, Spain; Department of Neurology, Hospital Universitari de Bellvitge, L’Hospitalet de Llobregat, Spain; Department of Neurology, Hospital Universitari Germans Trias i Pujol, Badalona, Spain; Genomic and Microscopy facilities, Germans Trias i Pujol Research Institute (IGTP), Badalona, Spain; Nephrology Service, Hospital Universitari Germans Trias i Pujol, Badalona, Spain

**Keywords:** Alzheimer disease, Dementia with Lewy bodies, miRNA, peripheral biomarkers, platelets, synucleinopathies

## Abstract

Dementia with Lewy bodies (DLB) is one of the most common causes of degenerative dementia after Alzheimer’s disease (AD) and presents pathological and clinical overlap with both AD and Parkinson’s disease (PD). Consequently, only one in three DLB cases is diagnosed correctly. Platelets, previously related to neurodegeneration, contain microRNAs (miRNAs) whose analysis may provide disease biomarkers. Here, we profiled the whole platelet miRNA transcriptome from DLB patients and healthy controls. Differentially expressed miRNAs were further validated in three consecutive studies from 2017 to 2019 enrolling 162 individuals, including DLB, AD, and PD patients, and healthy controls. Results comprised a 7-miRNA biosignature, showing the highest diagnostic potential for the differentiation between DLB and AD. Additionally, compared to controls, two miRNAs were down-regulated in DLB, four miRNAs were up-regulated in AD, and two miRNAs were down-regulated in PD. Predictive target analysis identified three disease-specific clusters of pathways as a result of platelet-miRNA deregulation. Our cross-sectional study assesses the identification of a novel, highly specific and sensitive platelet-associated miRNA-based bio-signature, which distinguishes DLB from AD.

**The paper explained:** *Problem:* Dementia with Lewy bodies (DLB) presents pathological and clinical overlap with both Alzheimer’s (AD) and Parkinson’s disease (PD), which impairs its correct diagnosis. Although numerous papers report peripheral biomarkers for AD, well-established biomarkers for DLB distinguishing it from AD are still missing. Platelet miRNA transcriptome was analyzed in several works, but their putative role as disease biomarkers for neurological disorders has not been assessed. It would be of paramount importance to establish a blood-based bio-signature as a minimally invasive mean for DLB diagnosis, improving differentiation of DLB patients from controls and AD.

*Results:* Our study revealed that platelet miRNAs might be promising biomarkers for the correct diagnosis of DLB stratifying patients in comparison with overlapping disorders, especially AD, and may help to highlight possible disease-related processes. In this cross-sectional study, which includes 162 individuals (DLB, AD, PD and healthy controls), platelet-associated miRNA content was disease group-specific. Three different miRNA sets together with their predicted targeted pathways were defined.

*Impact:* This study suggests that platelet miRNA may serve as DLB biomarker allowing the correct diagnosis and stratification in an easily-applied manner in clinical settings, and may help to highlight possible disease-related processes.

## Introduction

Dementia with Lewy bodies (DLB) is the second most common cause of degenerative dementia after Alzheimer’s disease (AD) and, together with Parkinson’s disease (PD), it belongs to the group of Lewy body disorders (LBD) (Ingelsson, 2016; Jellinger, 2018). Besides Lewy body pathology, a high percentage of DLB brains contains concomitant AD pathology (Colom-Cadena *et al*, 2017), also leading to the clinical overlap between DLB and AD. Although advances in the field have allowed improvements in their clinical characterization, it is still a challenge to diagnose DLB, AD and PD early and accurately (Jellinger, 2018). In particular, up to 80% of DLB cases are still misdiagnosed, usually as AD, and patients receive treatments that can adversely affect their cognition (McKeith *et al*, 2017), which makes the identification of biomarkers that permit the differential diagnosis of these diseases of paramount importance. Reduced Aβ42 levels have been found in AD-CSF (Ahmed *et al*, 2014) and tau and neurofilament are elevated in AD-plasma and CSF compared with controls (Zetterberg *et al*, 2013). Although classical AD CSF biomarkers have been explored as potential differential diagnosis tools in DLB patients, results from studies are controversial (Parnetti *et al*, 2019), and no peripheral biomarkers that differentiate between DLB and AD have been identified so far.

The study of blood and blood components has led to the identification of numerous circulating biomarkers. In particular, platelets are released into the circulation from the bone marrow after megakaryocytic differentiation, and although platelets are anucleate cells, they contain endoplasmic reticulum, ribosomes, and complete mitochondrial and apoptotic systems (Wojsiat *et al*, 2017). As well, a broad spectrum of functional mRNAs is found in platelets, which are translated after platelet activation. Since there are at least three different activation mechanisms, the resulting protein secretion profile depends on the specific activation pathway (Italiano *et al*, 2008; Milioli *et al*, 2015). Platelets are, therefore, able to modify their proteome in response to different environmental changes and stimuli (Leiter and Walker, 2019). Also, related to gene-expression regulation, the presence of miRNAs in platelets was described for the first time in 2008 (Bruchova *et al*, 2008) and, Landry and colleagues confirmed the existence of an almost complete and functional miRNA pathway one year later (Landry *et al*, 2009). Since then, several studies have been performed on the platelet miRNA content (Edelstein and Bray, 2011; Plé *et al*, 2012). In addition to their role in hemostasis and thrombosis, the functions of platelets include induction of apoptosis, initiation of immune response and tissue remodeling (Leiter and Walker, 2019). They have been described to show an enzymatic pathway similar to dopaminergic neurons (Wojsiat *et al*, 2017), and can store and release neurotransmitters, such as serotonin, glutamate and dopamine (Gowert *et al*. 2014). In AD, oxidative stress induces mitochondrial dysfunction and cell death in both platelets and neurons (Wojsiat *et al*, 2016; Zhao *et al*, 2016). Additionally, platelets contain up to 95% of the circulating form of the amyloid precursor protein (Zhao *et al*, 2016), they express several neuronal receptors and inflammatory-signalling molecules (Wojsiat *et al*, 2017), and also contain α-synuclein (Michell *et al*, 2005). Recently, it has also been shown that platelets play an active role during adult neurogenesis in the hippocampus (Seidemann *et al*, 2019). Finally, changes in activation, aggregation and morphology of platelets have been reported in PD, DLB, amyotrophic lateral sclerosis and multiple sclerosis (Behari and Shrivastava, 2013).

In this context, the aim of this study was to find out if platelet miRNAs may represent suitable biomarkers for DLB, distinguishing it from controls, and also from AD. First, we wanted to examine the complete platelet miRNA content in DLB compared to healthy individuals. Afterwards, to know if the detected differences were also detectable in independent cohorts and, if these profiles were disease-specific, we performed three independent validation studies, also including AD and PD patients. As a result, several sets of miRNAs differentiated DLB from the other groups of study subjects.

## Results

### Demographic and clinical data

Demographic and clinical data of patients are shown in **Table 1.** Mean age was similar between DLB patients, AD patients and controls, according to the inclusion criteria (75.1 +/- 6.8 years in the DLB group, 73.9 +/- 6.7 years in the AD group; p=0.236); however, PD patients were significantly younger (66.9 +/- 14.9 years, p=0.021). The male-female ratio was higher in PD and DLB, than in AD and CTRLs, but no gender-specific expression changes were observed during data analyses. Disease duration was similar between patients groups (p=0.068).

**Table 1:** Demographic and clinical data of the participants of the study.

|  | **DLB (n=59)** | **AD (n=28)** | **PD (n=24)** | **CTRLs (n=51)** | **p<sup>a</sup>** |
| --- | --- | --- | --- | --- | --- |
| Mean age, y<sup>b</sup> (age range, y) | 75.1 (57 – 89) | 73.9 (65 – 82) | 66.9 (42 – 87) | 72.0 (61 – 85) | 0.0067 |
| Gender (male/female ratio) | 1.5:1 | 1:1.5 | 1.5:1 | 1:1.5 | 0.493 |
| Disease duration<sup>c</sup>, y (range, y) | 4.2 (1.2 – 10.1) | 3.4 (0.8 – 5.6) | 5.2 (1.8 – 7.6) | - | 0.068 |
| MMSE<sup>d</sup>, mean (range) | 15.1 (3 – 28) | 20.3 (6 – 28) | - | - | 0.189 |
| UPDRS-III<sup>e</sup>, mean (range) | - | - | 20.9 (5 – 39) | - | - |
| GDS<sup>f</sup>, mean (range) | - | 4.1 (3 – 6) | - | - | - |
| Parkinsonism, n (%) | 47 (79.7%) | - | - | - | - |
| Positive DAT imaging, n (%) | 55 (93.2%) | - | - | - | - |

<sup>a</sup>p, obtained by the Kruskal-Wallis test; <sup>b</sup>y, years; <sup>c</sup>from the beginning of cognitive symptoms for DLB and AD, and motor symptoms in the case of PD; <sup>d</sup>MMSE, Mini-Mental Sate Examination; <sup>e</sup>UPDRs-III, Unified Parkinson's disease rating scale; <sup>f</sup>GDS, global deterioration scale.

### Platelet characterization and miRNA profile discovery

Analysis of the platelet-rich pellet for possible leukocyte contamination showed no staining for the leukocyte marker CD45 in our samples. Instead, we obtained a high fluorescent signal for the platelet marker CD61, indicating high platelet purity and no leukocyte contamination (**Figure 1A and Appendix Figure S1**).

**Figure 1.**
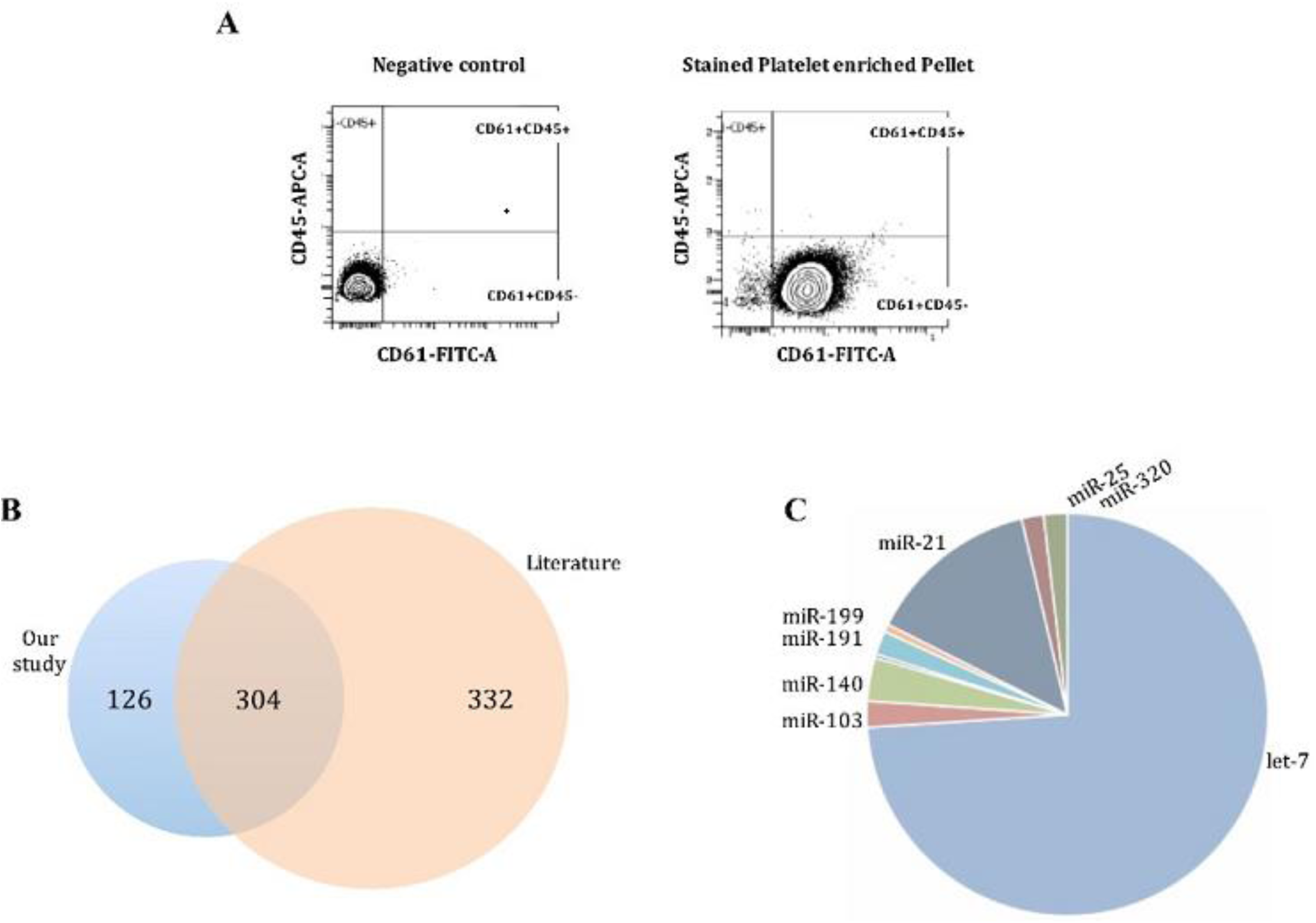
CD61+ platelet-rich pellet is enriched in previously described miRNAs. **A** CD61 staining was performed to identify platelets and CD45 was used as a leukocyte marker for staining of leukocyte contamination. Negative control with no CD61 staining (left); CD61-positive and CD45-negative staining is observed in the platelet-rich pellet (right). **B** Precursor-miRNA found in our study (blue circle) compared to literature (orange circle). **C** Most representative miRNA families found in our study.

RNA, used for the construction of NGS-libraries, showed an enriched profile of 20-40 nucleotide molecules characteristic for small RNA and miRNA. NGS generated a mean of 1,488,787±921,800 reads per sample in the control group, and 1,210,616±1,868,817 reads per DLB sample. These mapped to 1,279 known different mature miRNAs, and 534 miRNAs fulfilled the criteria of more than 5 reads per sample, corresponding to 430 different miRNA-precursor. Literature search revealed that 304 had been previously associated with platelets (**Figure 1B**), and 58.9% had been described in the first platelet-miRNA profiling studies (Landry *et al*, 2009; Osman *et al*, 2011). Our study also confirmed let-7, miR-103 and miR-21 (Plé *et al*, 2012) as the most common platelet-miRNA families (**Figure 1C**).

The normalized counts from NGS data were analyzed using the Wilcoxon-rank sum test, and 22 miRNAs were differentially expressed between DLB and controls, and were further validated by qPCR (hsa-miR-1343-3p, hsa-miR-191-3p, hsa-miR-6747-3p, hsa-miR-504-5p, hsa-miR-6741-3p, hsa-miR-128-3p, hsa-miR-1468-5p, hsa-miR-139-5p, hsa-let-7d-5p, hsa-let-7d-3p, hsa-miR-142-3p, hsa-miR-132-5p, hsa-miR-150-5p, hsa-miR-23a-5p, hsa-miR-26b-5p, hsa-miR-1301-3p, hsa-miR-625-3p, hsa-miR-146a-5p, hsa-miR-25-3p, hsa-miR-877-3p, hsa-miR-1908-5p, hsa-miR-744-5p).

### Validation of miRNA expression

The 22 differentially expressed miRNAs were validated by qPCR in three independent studies.

#### Study I (2017)

The first validation study included two cohorts of 21 DLB patients and 21 control individuals. Ten of the 22 miRNAs, hsa-miR-6747-3p, hsa-miR-128-3p, hsa-miR-139-5p, hsa-let-7d-5p, hsa-miR-142-3p, hsa-miR-132-5p, hsa-miR-150-5p, hsa-miR-26b-5p, hsa-miR-146a-5p, hsa-miR-25-3p, were diminished in DLB compared to controls (**Table 2**).

**Table 2:** Expression changes of the 22 miRNA identified as deregulated by NGS, in DLB versus controls. First validation study (2017).

| miRNA | expr change<sup>a</sup> | dev range<sup>b</sup> | p-value<sup>c</sup> |
| --- | --- | --- | --- |
| 1343-3p | 0.31 | 0.11-0.88 | 0.24 |
| 191-3p | 0.34 | 0.13-0.89 | 0.25 |
| **6747-3p** | **0.33** | **0.14-0.74** | **0.033** |
| 504-5p | 0.6 | 0.55-0.66 | 0.25 |
| 6741-3p | 0.32 | 0.16-0.66 | 0.17 |
| **128-3p** | **0.17** | **0.03-0.71** | **0.039** |
| 1468-5p | 0.37 | 0.16-0.89 | 0.28 |
| **139-5p** | **0.26** | **0.09-0.69** | **0.029** |
| **7d-5p** | **0.16** | **0.05-0.55** | **0.015** |
| 7d-3p | 0.34 | 0.12-0.95 | 0.21 |
| **142-3p** | **0.11** | **0.03-0.47** | **0.015** |
| **132-5p** | **0.21** | **0.06-0.70** | **0.018** |
| **150-5p** | **0.03** | **0.02-0.04** | **<0.0001** |
| 23a-5p | 0.58 | 0.41-0.81 | 0.31 |
| **26b-5p** | **0.16** | **0.04-0.62** | **0.017** |
| 1301-3p | 0.28 | 0.06-1.28 | 0.24 |
| 625-3p | 0.32 | 0.07-0.41 | 0.14 |
| **146a-5p** | **0.16** | **0.05-0.53** | **0.017** |
| **25-3p** | **0.20** | **0.06-0.66** | **0.034** |
| 877-3p | 0.34 | 0.14-0.84 | 0.22 |
| 1908-5p | 0.30 | 0.09-0.94 | 0.21 |
| 744-5p | 0.21 | 0.08-0.59 | 0.19 |

<sup>a</sup>expr change, expression change obtained by the deltadeltaCt method for DLB compared to controls; <sup>b</sup>dev range, deviation range of the expression change;
<sup>c</sup>p-value obtained by the Wilcoxon-Mann-Whitney followed by Dunn's test for multiple corrections. Significant expression changes are highlighted in bold.

#### Study II (2018)

Three independent cohorts comprising newly recruited DLB patients (n=22), AD patients (n=15) and control subjects (n=16) were included in the second validation study. Although 9 out of the ten miRNAs were again diminished in DLB when compared to controls, 5 out of these 9 miRNAs failed to produce significant results due to an elevated intra-group variability. Yet, four miRNAs confirmed the results of Study I. miRNAs hsa-miR-128-3p, hsa-miR-139-5p, hsa-miR-150-5p, hsa-miR-25-3p, were significantly down-regulated in DLB compared to controls, with hsa-miR-150-5p showing the most important decrease (**Table 3**).

**Table 3:**
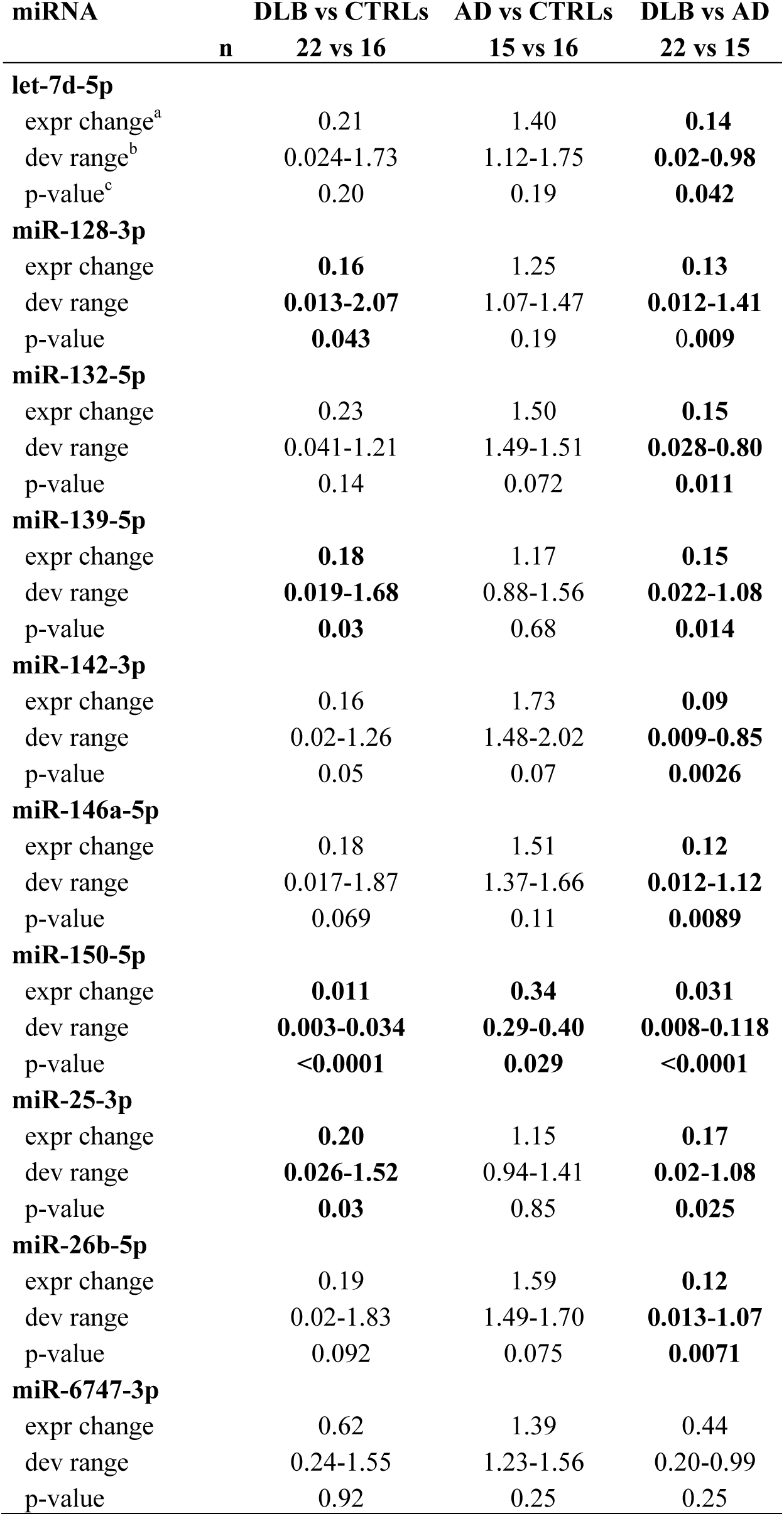
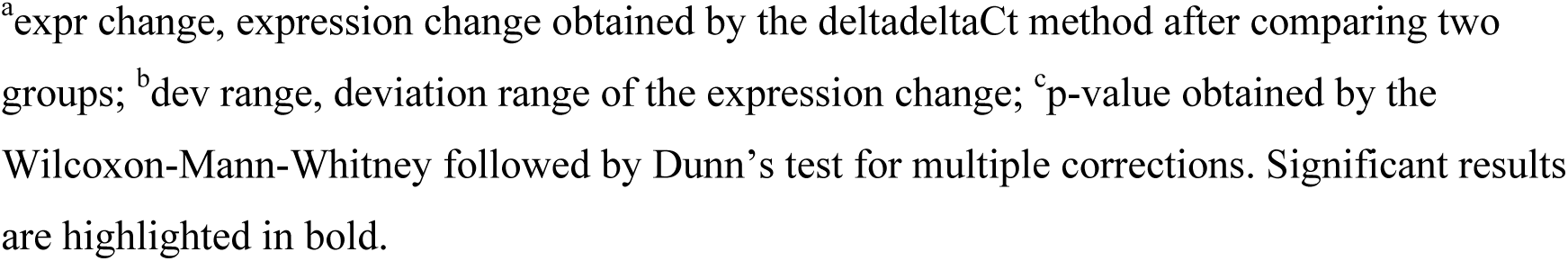
MiRNA expression results: Study II

| miRNA | DLB vs CTRLs | AD vs CTRLs | DLB vs AD |
| --- | --- | --- | --- |
| n | 22 vs 16 | 15 vs 16 | 22 vs 15 |
| **let-7d-5p** |  |  |  |
| expr change<sup>a</sup> | 0.21 | 1.40 | **0.14** |
| dev range<sup>b</sup> | 0.024-1.73 | 1.12-1.75 | **0.02-0.98** |
| p-value<sup>c</sup> | 0.20 | 0.19 | **0.042** |
| **miR-128-3p** |  |  |  |
| expr change | **0.16** | 1.25 | **0.13** |
| dev range | **0.013-2.07** | 1.07-1.47 | **0.012-1.41** |
| p-value | **0.043** | 0.19 | **0.009** |
| **miR-132-5p** |  |  |  |
| expr change | 0.23 | 1.50 | **0.15** |
| dev range | 0.041-1.21 | 1.49-1.51 | **0.028-0.80** |
| p-value | 0.14 | 0.072 | **0.011** |
| **miR-139-5p** |  |  |  |
| expr change | **0.18** | 1.17 | **0.15** |
| dev range | **0.019-1.68** | 0.88-1.56 | **0.022-1.08** |
| p-value | **0.03** | 0.68 | **0.014** |
| **miR-142-3p** |  |  |  |
| expr change | 0.16 | 1.73 | **0.09** |
| dev range | 0.02-1.26 | 1.48-2.02 | **0.009-0.85** |
| p-value | 0.05 | 0.07 | **0.0026** |
| **miR-146a-5p** |  |  |  |
| expr change | 0.18 | 1.51 | **0.12** |
| dev range | 0.017-1.87 | 1.37-1.66 | **0.012-1.12** |
| p-value | 0.069 | 0.11 | **0.0089** |
| **miR-150-5p** |  |  |  |
| expr change | **0.011** | **0.34** | **0.031** |
| dev range | **0.003-0.034** | **0.29-0.40** | **0.008-0.118** |
| p-value | **<0.0001** | **0.029** | **<0.0001** |
| **miR-25-3p** |  |  |  |
| expr change | **0.20** | 1.15 | **0.17** |
| dev range | **0.026-1.52** | 0.94-1.41 | **0.02-1.08** |
| p-value | **0.03** | 0.85 | **0.025** |
| **miR-26b-5p** |  |  |  |
| expr change | 0.19 | 1.59 | **0.12** |
| dev range | 0.02-1.83 | 1.49-1.70 | **0.013-1.07** |
| p-value | 0.092 | 0.075 | **0.0071** |
| **miR-6747-3p** |  |  |  |
| expr change | 0.62 | 1.39 | 0.44 |
| dev range | 0.24-1.55 | 1.23-1.56 | 0.20-0.99 |
| p-value | 0.92 | 0.25 | 0.25 |

<sup>a</sup>expr change, expression change obtained by the deltadeltaCt method after comparing two groups; <sup>b</sup>dev range, deviation range of the expression change; <sup>c</sup>p-value obtained by the Wilcoxon-Mann-Whitney followed by Dunn's test for multiple corrections. Significant results are highlighted in bold.

The comparison of DLB and AD miRNA expression data revealed that 9 out of the 10 miRNAs were significantly down-regulated in DLB compared with AD (**Table 3**). Only hsa-miR-150-5p was significantly diminished in AD when compared to controls.

#### Study III (2019)

To the initially recruited patients, four independent cohorts of newly diagnosed DLB (n=16), AD (n=13) and PD patients (n=24), and 14 control subjects were added in the third validation study, analyzing a total of 162 individuals (59 DLB patients, 28 AD patients, 24 PD patients, 51 controls). As a result, two miRNAs (hsa-miR-142-3p, hsa-miR-150-5p) were significantly diminished in DLB compared with controls. Seven miRNAs (hsa-let-7d-5p, hsa-miR-142-3p, hsa-miR-132-5p, hsa-miR-150-5p, hsa-miR-26b-5p, hsa-miR-146a-5p, hsa-miR-25-3p,) were significantly diminished in DLB compared to AD, and two (hsa-miR-150-5p and hsa-miR-26b-5p) were down-regulated in DLB compared to PD (**Table 4**). When grouping DLB and PD as LBD, only hsa-miR-139-5p was significantly down-regulated, but hsa-miR-128-3p and hsa-miR-139-5p were diminished in PD compared to controls. The expression of four miRNAs (hsa-miR-132-5p, hsa-miR-146a-5p, hsa-miR-25-3p, hsa-miR-6747-3p) was elevated in AD *vs* CTRLs (**Table 4, Figure 2**). Of all miRNAs, only hsa-miR-150-5p was differentially expressed in DLB compared to each of the other cohorts, suggesting disease-specific deregulation (**Appendix Figure S2**).

**Table 4:**
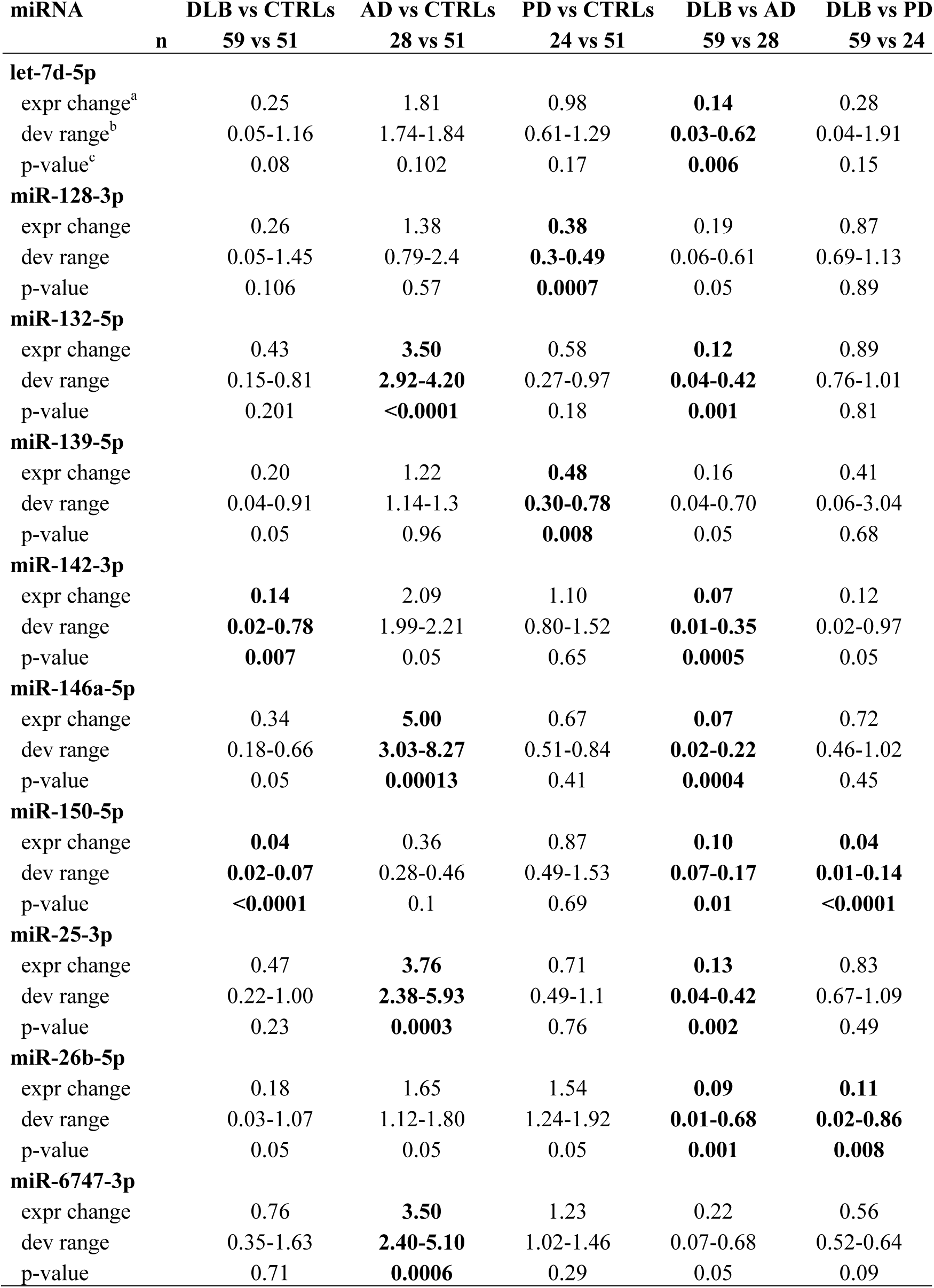
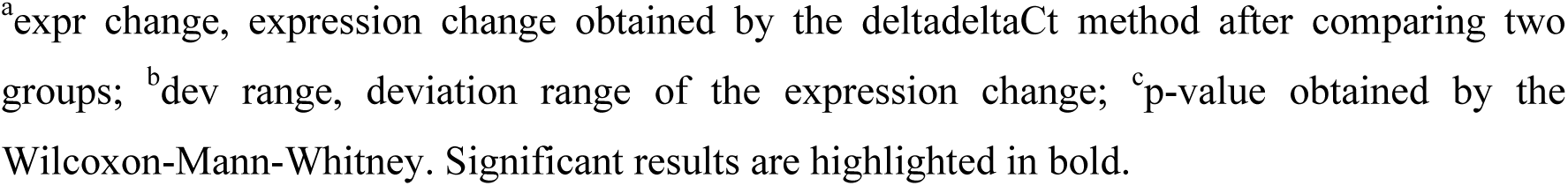
Combined results of Studies I-III.

| miRNA | DLB vs CTRLs | AD vs CTRLs | PD vs CTRLs | DLB vs AD | DLB vs PD |
| --- | --- | --- | --- | --- | --- |
| n | 59 vs 51 | 28 vs 51 | 24 vs 51 | 59 vs 28 | 59 vs 24 |
| **let-7d-5p** |  |  |  |  |  |
| expr change<sup>a</sup> | 0.25 | 1.81 | 0.98 | **0.14** | 0.28 |
| dev range<sup>b</sup> | 0.05-1.16 | 1.74-1.84 | 0.61-1.29 | **0.03-0.62** | 0.04-1.91 |
| p-value<sup>c</sup> | 0.08 | 0.102 | 0.17 | **0.006** | 0.15 |
| **miR-128-3p** |  |  |  |  |  |
| expr change | 0.26 | 1.38 | **0.38** | 0.19 | 0.87 |
| dev range | 0.05-1.45 | 0.79-2.4 | **0.3-0.49** | 0.06-0.61 | 0.69-1.13 |
| p-value | 0.106 | 0.57 | **0.0007** | 0.05 | 0.89 |
| **miR-132-5p** |  |  |  |  |  |
| expr change | 0.43 | **3.50** | 0.58 | **0.12** | 0.89 |
| dev range | 0.15-0.81 | **2.92-4.20** | 0.27-0.97 | **0.04-0.42** | 0.76-1.01 |
| p-value | 0.201 | **<0.0001** | 0.18 | **0.001** | 0.81 |
| **miR-139-5p** |  |  |  |  |  |
| expr change | 0.20 | 1.22 | **0.48** | 0.16 | 0.41 |
| dev range | 0.04-0.91 | 1.14-1.3 | **0.30-0.78** | 0.04-0.70 | 0.06-3.04 |
| p-value | 0.05 | 0.96 | **0.008** | 0.05 | 0.68 |
| **miR-142-3p** |  |  |  |  |  |
| expr change | **0.14** | 2.09 | 1.10 | **0.07** | 0.12 |
| dev range | **0.02-0.78** | 1.99-2.21 | 0.80-1.52 | **0.01-0.35** | 0.02-0.97 |
| p-value | **0.007** | 0.05 | 0.65 | **0.0005** | 0.05 |
| **miR-146a-5p** |  |  |  |  |  |
| expr change | 0.34 | **5.00** | 0.67 | **0.07** | 0.72 |
| dev range | 0.18-0.66 | **3.03-8.27** | 0.51-0.84 | **0.02-0.22** | 0.46-1.02 |
| p-value | 0.05 | **0.00013** | 0.41 | **0.0004** | 0.45 |
| **miR-150-5p** |  |  |  |  |  |
| expr change | **0.04** | 0.36 | 0.87 | **0.10** | **0.04** |
| dev range | **0.02-0.07** | 0.28-0.46 | 0.49-1.53 | **0.07-0.17** | **0.01-0.14** |
| p-value | **<0.0001** | 0.1 | 0.69 | **0.01** | **<0.0001** |
| **miR-25-3p** |  |  |  |  |  |
| expr change | 0.47 | **3.76** | 0.71 | **0.13** | 0.83 |
| dev range | 0.22-1.00 | **2.38-5.93** | 0.49-1.1 | **0.04-0.42** | 0.67-1.09 |
| p-value | 0.23 | **0.0003** | 0.76 | **0.002** | 0.49 |
| **miR-26b-5p** |  |  |  |  |  |
| expr change | 0.18 | 1.65 | 1.54 | **0.09** | **0.11** |
| dev range | 0.03-1.07 | 1.12-1.80 | 1.24-1.92 | **0.01-0.68** | **0.02-0.86** |
| p-value | 0.05 | 0.05 | 0.05 | **0.001** | **0.008** |
| **miR-6747-3p** |  |  |  |  |  |
| expr change | 0.76 | **3.50** | 1.23 | 0.22 | 0.56 |
| dev range | 0.35-1.63 | **2.40-5.10** | 1.02-1.46 | 0.07-0.68 | 0.52-0.64 |
| p-value | 0.71 | **0.0006** | 0.29 | 0.05 | 0.09 |

<sup>a</sup>expr change, expression change obtained by the deltadeltaCt method after comparing two groups; <sup>b</sup>dev range, deviation range of the expression change; <sup>c</sup>p-value obtained by the Wilcoxon-Mann-Whitney. Significant results are highlighted in bold.

**Figure 2.**
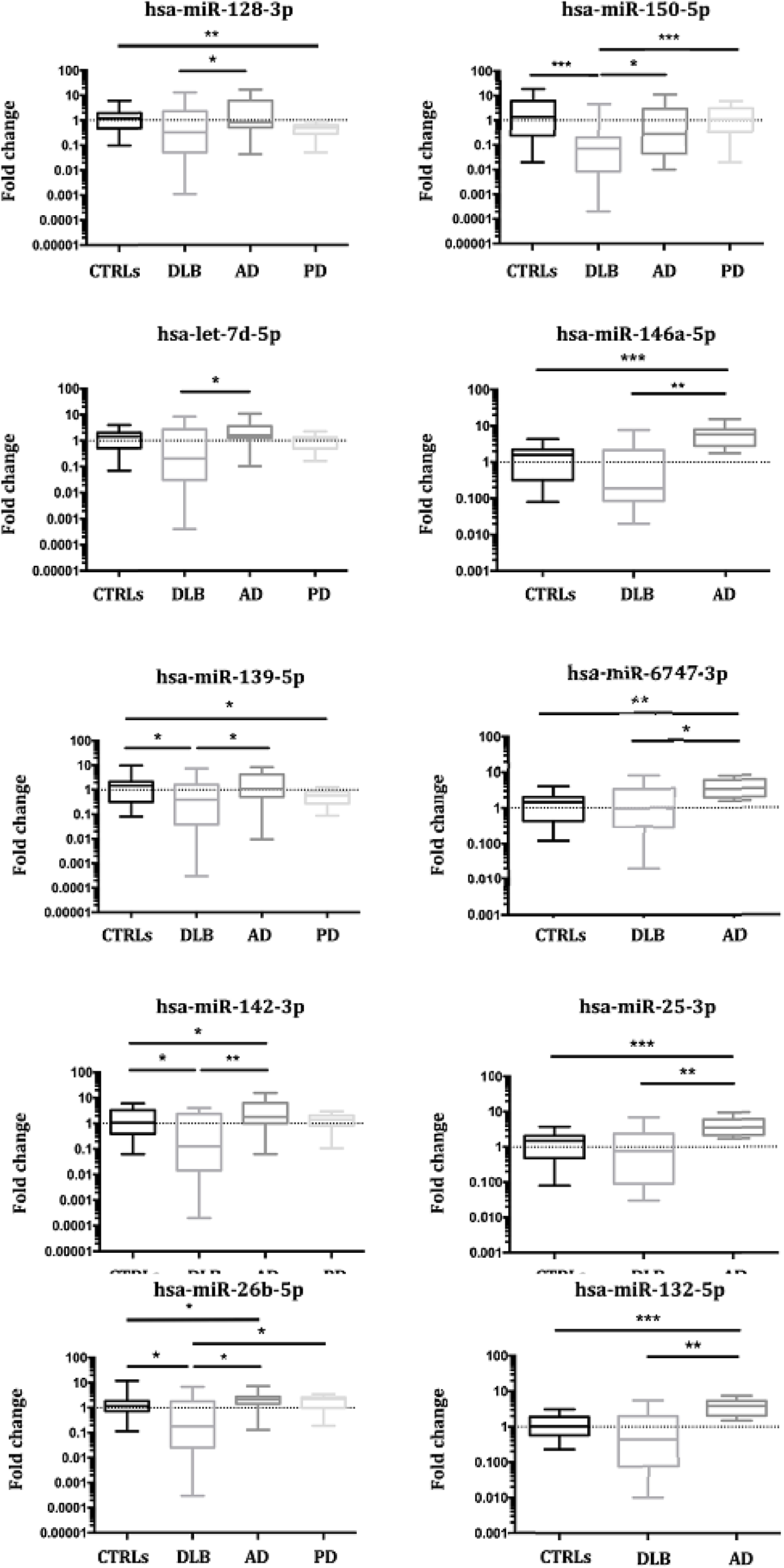
Combined results from three validation studies for miRNA expression in Controls, DLB, AD and PD. In all cases, mean and range for fold change are plotted; (*p<0.05, **p<0.001, ***p<0.0001).

The 5 miRNA sets were further studied for their usefulness as biomarkers (**Figure 3A**).

**Figure 3.**
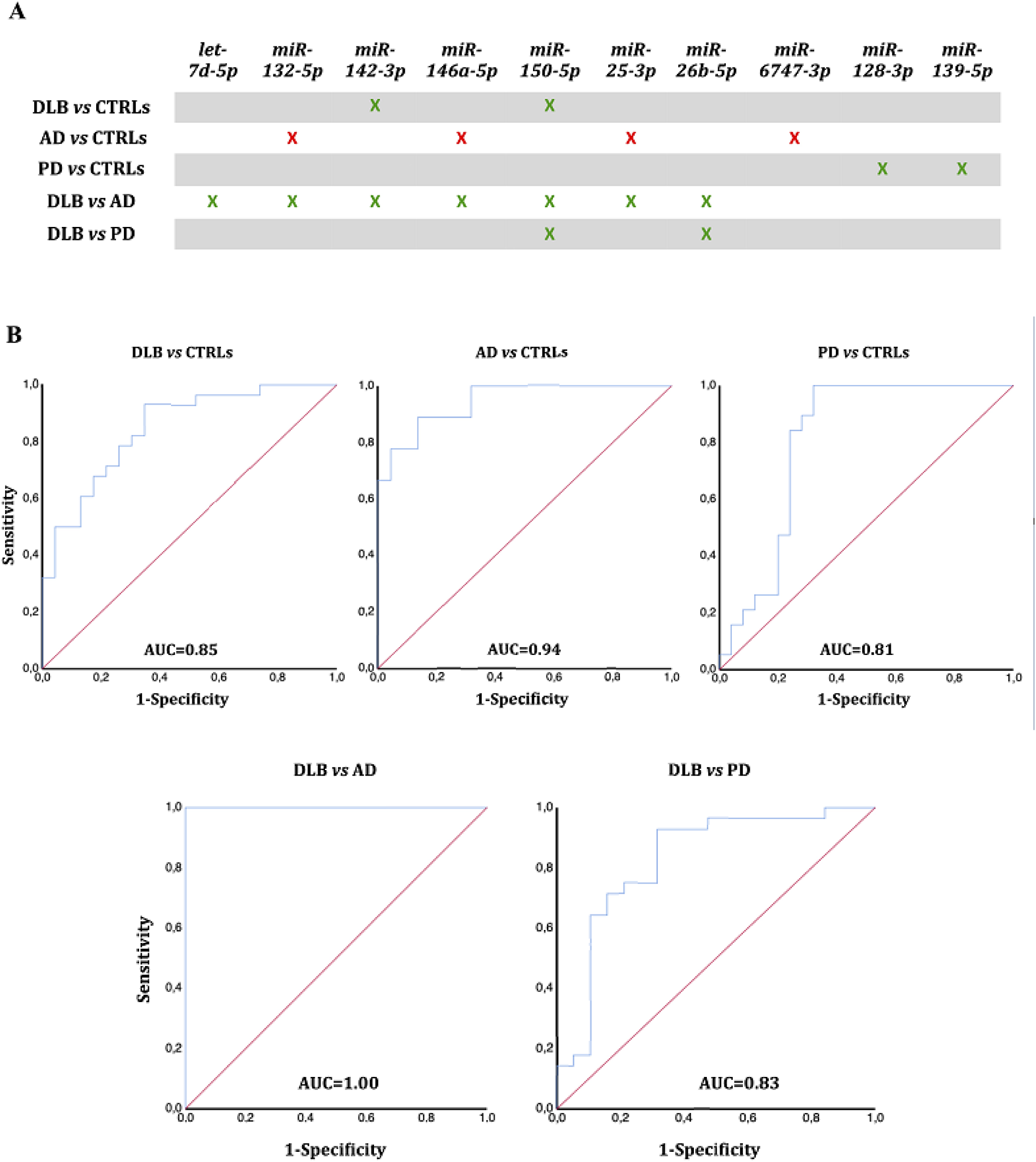
Diagnostic outcome for the five miRNA-sets. **A** Significantly different miRNAs (p<0.01) were clustered in 5 different sets. Diminished expression – green, increased expression – red. **B** ROC curves were calculated for miRNAs with differential expression (p<0.01) between two cohorts.

### ROC curve analysis

ROC curves were calculated for all five miRNA sets to assess their discrimination potential between groups. The combination of the seven differentially expressed miRNAs between DLB and AD (miRNAs hsa-let-7d-5p, hsa-miR-132-5p, hsa-miR-142-3p, hsa-miR-146a-5p, hsa-miR-150-5p, hsa-miR-25-3p and hsa-miR-26b-5p) presented the highest specificity (100%) and sensitivity (100%) to distinguish DLB patients from AD patients, with an AUC of 1 (**Figure 3B**).

The ROC curve for hsa-miR-142-3p and hsa-miR-150-5p, differentially expressed between DLB and CTRLs, yielded an AUC=0.85 (95% C.I. 0.74-0.95; 82% sensitivity, 70% specificity). Comparison of AD and CTRLs, miRNAs hsa-miR-132-5p, hsa-miR-146a-5p, hsa-miR-25-3p, and hsa-miR-6747-3p resulted in AUC=0.94 (95% C.I. 0.86-1.00; 89% sensitivity, 80% specificity); and AUC=0.81 (95% C.I. 067-0.94; 84% sensitivity, 76% specificity) was obtained for hsa-miR-128-3p and hsa-miR-139-5p comparing PD and CTRLs. AUC=0.83 (95% C.I. 0.73-0.98; 90% sensitivity, 73.7% specificity), was obtained for hsa-miR-150-5p and hsa-miR-26b-5p when comparing DLB and PD (**Figure 3B**).

### miRNA expression in whole blood

To assess whether the results were platelet-specific, we analyzed the 10 differentially expressed miRNAs in whole blood of DLB, PD and AD patients, and controls (n=16, each). Only the expression of hsa-let-7d-5p and hsa-miR-132-5p was diminished in blood of PD patients in comparison with controls (**Appendix Table S1**). Specifically, these miRNAs did not show expression changes in platelets of PD patients. No additional differences were found.

### miRNA target prediction

To obtain a list of possible target genes, miRTarbase (Chou *et al*, 2018) and miRGate (Andrés-León *et al*, 2017) were used to perform predictive analyses for the miRNAs sets with differential expression between DLB and CTRLs, DLB and AD, DLB and PD, AD and CTRLs, and PD and CTRLs.

When comparing DLB and controls, the target screening of miRNAs hsa-miR-150-5p and hsa-miR-142-3p identified 81 different genes. Genes involved in transcriptional regulation (p=0.001), cellular response to stress (p=8.36·10^−5^) and immune response (p=0.021), were overrepresented (**Figure 4A**).

**Figure 4.**
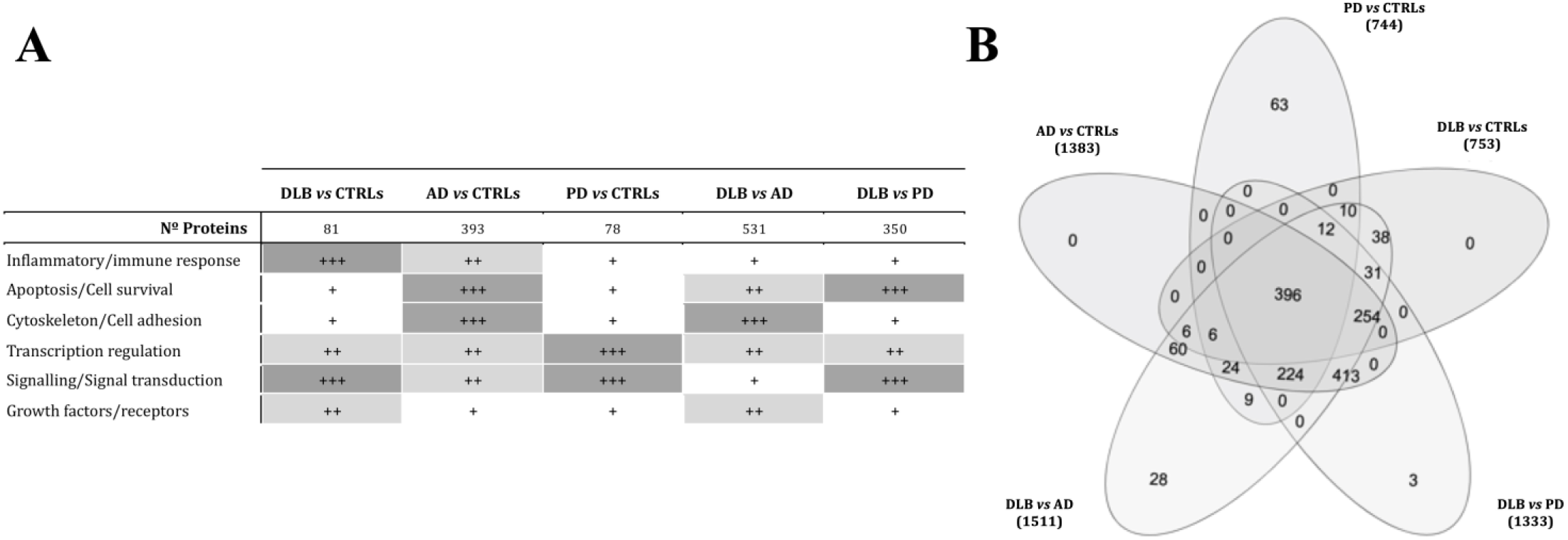
Target prediction analysis for the 5 miRNA sets was performed with miRTarbase and miRGate online tools. **A** Most relevant targeted pathways, according to Reactome and String analysis for each target-gene list. Mostly, genes related to transcriptional regulation and signal transduction were overrepresented (+++, ≥40% of the pathways related; ++, 20-40% of related pathways; +, up to 20% of the defined pathways). **B** Venn diagram comparing altered pathways in the 5 miRNA-sets defined. In PD, 63 pathways were specifically altered, and 31 were associated with DLB in comparison to AD and PD.

The seven miRNAs down-regulated in DLB compared to AD were predicted to target genes involved in integrin cell surface interactions (p=1.5·10^−4^), cell death pathways (p=0.002), and transcriptional regulation (p=0.004) (**Figure 4A**). When analyzing miRNAs diminished in DLB compared with PD, formation of senescence-associated heterochromatin foci (SAHF) was the most representative pathway (p=2.4·10^−4^).

The analysis of the miRNA-set that distinguishes AD from controls (**Figure 4A**) rendered 393 possible target genes. Of these, 27.7% were involved in transcriptional regulation, 43.3% in signal transduction and 54.2% were phosphoprotein-coding. Gene clusters related to stress (p=8.3·10^−5^) and immune response (p=1.58·10^−5^) were identified, including *TDRD7* (tudor-domain-containing protein 7) and *TIA1* (TIA1 cytotoxic granule associated RNA binding protein), both involved in stress granule formation.

MiRNAs hsa-miR-128-3p and hsa-miR-139-5p, down-regulated in PD compared to controls, were predicted to target 78 different genes. Signal transduction (p=1.38·10^−10^) and PIP-AKT signalling (p=4.9·10^−10^) were the most enriched pathways (**Figure 4A**). A protein-phosphorylation (p=0.0376) and a MAPK-pathway (p=0.015) cluster were identified including *FOS, MTOR* and *RICTOR* (**Figure 4A)**.

Comparison of the five pathway lists revealed that 31 pathways were altered specifically in DLB (**Figure 4B**). RNA and small RNA metabolism (mitochondrial tRNA processing), and RNA silencing by small RNA were found. In AD, specific pathway enrichment comprised cell death-related pathways (p=5.04·10^−8^). In PD, 63 pathways were specifically enriched (**Figure 4B**), including MAP kinase, protein phosphorylation pathways (p=8.3·10^−8^), negative regulation of cell death (p=0.009), and serotonin and dopamine receptor-related pathways (p=0.05).

## Discussion

In this study, we analysed the platelet miRNA profile in DLB patients, compared with AD and PD patients, and also with healthy controls, aiming to identify biomarkers for DLB. As a result of the first NGS discovery phase, we selected 22 differentially expressed miRNAs which were further validated in three independent qPCR-based studies, including independent cohorts of DLB, AD, PD, and controls. Since the clinical diagnosis of DLB is still challenging, primarily because of its overlap with AD but also with PD (Mckeith *et al*, 2017), there is an urgent need for biomarkers to differentiate between these neurodegenerative disorders. Here, we defined 3 different groups of miRNAs as specifically deregulated in each of the three neurodegenerative disorders.

The first group was DLB-specific, consisted of 7 miRNAs and comprised three subsets. Hsa-miR-142-3p and hsa-miR-150-5p showed diminished expression compared to controls; these two miRNAs, together with hsa-let-7d-5p, hsa-miR-132-5p, hsa-miR-146a-5p, hsa-miR-25-3p and hsa-miR-26b-5p were down-regulated compared to AD, and hsa-miR-150-5p and hsa-miR-26b-5p were decreased in comparison with PD. Putative target genes were related to cell senescence, inflammation and signalling, and to RNA metabolism, especially to mitochondrial tRNA and gene silencing by small RNA. The disruption of RNA metabolism alterations in RNA splicing and processing, together with the deregulation of non-coding RNA has been described in several brain disorders (Nussbacher *et al*, 2019). In early PD brains, alterations in the small RNA profile are specifically related to tRNA fragments (Pantano *et al*, 2015). However, the relation between the impairment of these pathways and the development of DLB remains to be determined.

The second group, hsa-miR-132-5p, hsa-miR-146a-5p and hsa-miR-6747-3p, was up-regulated in AD. Among the predicted target genes, we found apoptosis and cell-death, together with stress response-related genes. Specifically, *TIA1*, required for the formation of stress granules (Apicco *et al*, 2018), and *TDRD7*, a component of cytoplasmic RNA granules (Lachke *et al*, 2011), were identified. *TIA1* is involved in RNA splicing and post-transcriptional gene regulation, has been found in stress granules (Wolozin and Ivanov, 2019), and has been related with the tau oligomer and neurofibrillary tangle deposition pattern in AD (Apicco *et al*, 2018). Stress granules are formed in the cytoplasm during transient cellular stress, and their nature and biology could be altered in neurodegenerative diseases, with chronic and long-term stress (Wolozin and Ivanov, 2019).

The third group of miRNAs, hsa-miR-128-3p and hsa-miR-139-5p, were significantly decreased in PD. Although transcriptional regulation and signal transduction were enriched in all miRNA sets, these were importantly over-represented among the target genes for these two PD-specific miRNAs. Both *RICTOR* and *MTOR* are predicted targets for hsa-miR-128-3p and play an essential role in neuronal survival and synaptic plasticity (Pérez-Santiago *et al*, 2019). In PD brains, MTOR expression and AKT functions are impaired (Martín-Flores *et al*, 2018). MTOR over-expression impairs autophagy in genetic PD, enhancing α-synuclein deposition (Zhu *et al*, 2019), and hsa-miR-128-3p down-regulation could be involved in the up-regulation of this pathway.

To our knowledge, the 7-miRNA biosignature composed of hsa-miR-142-3p, hsa-miR-150-5p, hsa-let-7d-5p, hsa-miR-132-5p, hsa-miR-146a-5p, hsa-miR-25-3p and hsa-miR-26b-5p, represents the first molecular signature that permits to distinguish DLB from AD with high specificity and sensitivity. Although the precise involvement of these miRNAs in DLB pathology has yet to be clarified, the identification of these biomarker candidates is particularly important, because they may improve DLB diagnosis and correspondingly, patient management, treatment and outcome. The 4^th^ consensus report of the DLB Consortium underlined the urgent need of developing guidelines and outcome measures for clinical trials in DLB (McKeith *et al*, 2017), this study could be crucial in the identification of a diagnostic biomarker to define inclusion/exclusion criteria for either patients with DLB, PD or AD in clinical trials.

Interestingly, three disease-specific clusters of pathways and biological processes were identified as the result of platelet-miRNA deregulation. In DLB, pathways were related to gene expression and small RNA metabolism; in AD, to stress response; and in PD, to protein phosphorylation, metabolism and degradation. Since DLB and PD are synucleinopathies, the identification of rather similar pathways could have been expected. However, since none of the PD patients had developed dementia when samples were obtained, the involvement of different pathways may reflect the mechanisms leading to early dementia development in synucleinopathies. The study of PD patients with dementia is needed to elucidate this question further.

No differences in miRNA expression were found in whole blood, indicating that platelet-specific miRNA deregulation could be related to disease pathogenesis. Since platelets present neuron-like metabolic pathways, previous studies have shown that in AD, APP acts as a platelet-membrane receptor contributing mostly to soluble β-amyloid after platelet activation (Bram *et al*, 2018). Mitochondrial dysfunction, higher content in phosphorylated TDP43, and morphological and structural platelet changes in AD and PD have also been reported (Koçer *et al*, 2013; Kucheryavykh *et al*, 2017). Whether miRNA deregulation in platelets promotes neurodegeneration or merely reflects its effects remains to be elucidated. But a possible link between platelets and brain plasticity has been recently described (Leiter and Walker, 2019), showing that platelets act directly on neural precursor cells *in vitro*, and that specific exercise-induced platelet activation leads to enhanced hippocampal neurogenesis (Leiter *et al*, 2019).

Although this study has been performed in a multi-centre setting, our results need to be replicated by independent studies in other laboratories and in multi-national studies with larger cohorts. Further research should also address and confirm the alteration of the predicted biological pathways and their relationship with DLB, AD and PD. Additionally, these miRNAs should also be analyzed in groups of individuals at risk of developing a synucleinopathy or dementia, as could be individuals with idiopathic REM sleep behaviour disorder (IRBD) and mild cognitive impairment (MCI).

In summary, two main findings must be highlighted. First, we have shown that the miRNA content from platelets may represent a promising source of biomarkers. In particular, we defined a 7-miRNA bio-signature that may represent a useful biomarker for the differentiation between DLB and AD patients. Second, we defined specific clusters of pathways and biological processes for DLB, AD and PD, underlining that the development of the different diseases is, at least in part, platelet-driven by affecting specific pathways.

## Materials and methods

The whole workflow of this study is shown in **Appendix Figure S3.**

### Participants

The current study was conducted between 2015 and 2019. A total of 162 individuals were included from two different hospitals: Hospital Universitari Germans Trias i Pujol (Badalona, Barcelona) and Hospital Universitari de Bellvitge (L’Hospitalet de Llobregat, Barcelona). Four cohorts were recruited:

#### DLB patients

Fifty-nine patients who fulfilled criteria for probable DLB (McKeith *et al*, 2005; 2017) were prospectively recruited from those visited in the Neurodegenerative disease Unit of both centres as routine clinical practice.

#### AD patients

Twenty-eight patients who fulfilled criteria for probable AD (National Institute on Aging-Alzheimer’s Association criteria, NIA-AA 2011; McKhann *et al*, 2011) were also consecutively recruited at the Neurodegenerative disease Unit at their routine visits, irrespective of any specific complaint or clinical feature. AD patients were matched by age with the DLB patients.

#### PD patients

For comparison purposes, a group of 24 PD patients diagnosed according to the UK PD Society Brain Bank criteria (Hughes *et al*, 1992) were included. None of these patients presented cognitive impairment, which was defined as subjective cognitive complaints, based on the patient’s and informant interview, and on the Minimental State Examination (MMSE) score, considering cognitive impairment if the MMSE punctuation was < 24 points.

In DLB, AD and PD patients, age at onset was defined as the age when memory loss or parkinsonism was first noticed by the patient or his/her relatives.

#### Control subjects

Fifty-one control individuals were selected among non-blood relatives of the patients, age-matched with the DLB group.

The study was carried out in three independent phases; the first in 2017 included 21 DLB patients and 21 controls, the second in 2018 comprised 22 DLB, and 15 AD patients, and 16 controls, and the third in 2019 contained 16 DLB, 13 AD and 24 PD patients, and 14 controls.

The study was carried out with the approval of the local Ethical Committees for Clinical Investigation of the institutions involved in the study, and a written informed consent was signed by all participants or their legal guardians according to the Declaration of Helsinki (Lynöe *et al*, 1991).

### Blood collection, purification and characterization of platelets

Peripheral blood was collected following standard procedures to minimize coagulation and platelet activation (György *et al*, 2014). After venous puncture, blood was collected in sodium citrate tubes (BD Vacutainer®, New Jersey, USA), and processed within 2 hours following collection. After centrifugation at 500 x g for 10 minutes at room temperature to pellet red blood cells and leukocytes, the supernatant was centrifuged at 2,500 x g for 15 minutes at room temperature to obtain a platelet-rich pellet (Sáenz-Cuesta *et al*, 2015). The pellet was re-suspended in 250 µL of PBS and characterized by flow cytometry for sample purity according to typical platelet size and complexity (FSC/SSC) using 100 um-Red Nile Beads (ThermoFisher) as reference and phenotypically confirmed as CD45-/CD61+ (ImmunoTools, ref21270456 and ref21330613, respectively). The analysis was performed on a FACSCanto II flow cytometer (BD).

The samples were stored at -80°C until further processing.

### Purification of platelet-derived small RNA

Platelet-rich pellets were thawed on ice. miRNA isolation was performed using the mirVana Paris Kit (Invitrogen). Briefly, 600 µL of lysis buffer and 1/10 of miRNA Homogenate Additive Mix were added to each pellet and incubated for 10 minutes on ice after vortexing. One volume of phenol-chloroform was added, mixed and centrifuged at 10,000 x g for 5 minutes. One-third and two-thirds volume of ethanol was added in 2 consecutive steps to the miRNA containing aqueous phase and passed through a filter column. After the recommended washing steps, miRNAs were obtained with 100 µL of elution buffer and stored at -80°C until later analysis.

### MiRNA isolation from whole blood

RNA isolation was carried out after collection of 3 ml of whole blood in PAXgene Blood RNA tubes (PreAnalytiX, Hombrechtikon, Switzerland) with the PAXgene Blood miRNA Kit 50, v2 (PreAnalytiX) following manufacturer’s instructions. RNA quantity, purity and integrity were ascertained by the Agilent 2100 Bioanalyzer (Agilent Technologies, Santa Clara, USA).

### Discovery Phase: miRNA sequencing and sequencing data analysis

The total miRNA volume obtained from 7 DLB and 7 control samples was precipitated overnight at -20°C with 1 µL of glycogen (20 µg/ µL), 10% 3 M AcNa (ph 4.8) and 2 volumes of ethanol. miRNAs were resuspended in 10 µL RNase free H_2_O and heated at 65°C for 3 min. Quality control and size distribution of the purified small RNA was assessed by Bioanalyzer 2100 (Agilent Technologies).

Six µL of each sample were used for library preparation with the NEBNext Multiplex Small RNA Sample Preparation Set for Illumina (New England Biolabs) following the manufacturer’s instructions. Individual libraries were subjected to quality analysis using a D1000 ScreenTape (TapeStation, Agilent Technologies), quantified by fluorometry and pooled. Clustering and sequencing were performed on an Illumina Sequencer (MiSeq, Illumina, San Diego, USA) at 1 x 50c single read mode, and 200,000 reads were obtained for each sample.

### Validation Phase: Reverse transcription and quantitative real-time PCR

MiRNA was reverse-transcribed using the MiRCURY LNA™ Universal cDNA Synthesis Kit II (Exiqon, Vedbaek, Denmark) according to the manufacturer’s protocol. After adjusting RNA concentration to 5 ng/µL and mixing with reaction buffer and enzyme mix, a retro-transcription reaction was carried out at 42°C for 60 minutes. Artificial RNA UniSp6 from the same kit was used as a retro-transcription control. Quantitative PCR (qPCR) was performed on a LightCycler 480 (Roche, Basel, Switzerland) using miRNA LNA technology and Pick&Mix PCR pre-designed panels (Exiqon) with miRNA UniSp3 as interplate calibrator control. cDNA was diluted 1:80, 4 µL were used with ExiLENT SYBR Green Master Mix (Exiqon, Vedbaek, Denmark) following manufacturer’s indications and samples were set up in duplicate.

The validation study was carried out in three independent phases, the first in 2017, the second in 2018 and the third in 2019, including the subjects as described in 2.1.

### Statistical analysis

Values for NGS data and reads are given as mean ± SD. Expression levels of the miRNAs selected for qPCR validation were determined using crossing point (Cp) values. Cp values were averaged between duplicates and normalized against UniSp6 spike-in Cp values for platelet-derived miRNA and against hsa-miR-191-5p in the case of whole blood. Relative expression changes were calculated by the -ΔΔCt method (Schmittgen and Livak, 2008) and the results were further evaluated with the Wilcoxon-Mann-Whitney test (https://ccb-compute2.cs.uni-saarland.de/wtest/) and the two-tailed unpaired T-test to compare the expression between two groups. When comparing more than two groups (DLB, controls, AD and PD), multiple comparisons were performed using the Kruskal-Wallis non-parametric test and Dunn’s test was used for multiple corrections (GraphPad Software, Inc., La Jolla, CA, USA). In all cases, a confidence interval of 95% and a p-value below 0.05 was considered to be significant. To assess the diagnostic potential, the area under the ROC curve (AUC) was calculated for miRNAs with p-value < 0.01 by the Wilson/Brown method using SPSS Statistics 21 (IBM, Armonk, NY, USA) and GraphPadPrism v7 in order to determine the diagnostic sensitivity and specificity (95% C.I., AUC > 0.80 was considered as the minimum value for a useful biomarker).

### miRNA target prediction and analysis

Possible targets of deregulated miRNAs (those, showing p-value <0.01) were predicted using miRTarbase (Chou *et al*, 2018) accepting as target genes those that were reported only by strong evidence studies; and by miRGate (Andrés-León *et al*, 2017), considering only confirmed targets. For each miRNA set, including miRNAs with expression change of miRNAs of p < 0.01, and identified as disease-specific, targets from both databases were taken together, and overlapping data were removed before screening the complete list for their molecular relationship with String DataBase (https://string-db.org/; Szklarczyk *et al*, 2017) and the Reactome online tool (Fabregat *et al*, 2018). Gene description and most relevant information were screened also through the Uniprot database (Pundir *et al*, 2017). For each miRNA set, target genes were clusters by their functional characteristics, and related molecular pathway.

### Data availability

The RNA-seq data were deposited in NCBI GEO database (https://www.ncbi.nlm.nih.gov/geo/, series GSE147218) and at SRA (https://www.ncbi.nlm.nih.gov/sra/, BioProject-ID: PRJNA613191).

## Acknowledgements

We thank all participants for providing their blood samples for this study, together with the members of the Department of Neurology from both hospitals, and Àngels Barberà Pla from the Pathology Department of Germans Trias i Pujol Research Institute, who made the validation of this study possible. We also thank Anna Oliveira from the Genomics Unit (Health Sciences Research Institute Germans Trias i Pujol), Dr Mireia Coma from ANAXOMICS Biotech S.L. (Barcelona) and Dr Sonia Jansa (BioNova Científica, S.L.) for their support and guiding for genomic data processing and analysis.

## Author contributions

AG-V, FEB and KB conducted study design. AG-V performed experiments, data acquisition, and analyses. JC, DV, LI, DS, JG-B, RR, RA stratified patients, collected and analysed clinical samples and manage, together with KB, patients’ information. MPA designed and guided RNA and sequencing experiments and analysis. AG-V, FEB and KB wrote the manuscript, reviewed and agreed by all coauthors.

## Legends of Appendix Supplementary Figures

**Appendix SFigure 1. The complete workflow of the current study.** A first discovery phase by Next-generation sequencing included a cohort of 7 DLB patients and 7 healthy controls. Selected miRNAs were validated in three independent qPCR validation studies.

**Appendix SFigure 2. Additional samples analyzed by flow cytometry for the** expression of CD61 as platelet marker and CD45 as leukocyte marker.

**Appendix SFigure 3. hsa-miR-150-5p expression in DLB, AD, PD and controls.**

**A** Expression change by qPCR. Mean, and Min to Max values are plotted.

**B-D** ROC curves for hsa-miR-150-5p as biomarker for DLB cases *vs* controls (B), for DLB *vs* PD (C) and for DLB *vs* AD (D).

